# A confining microfluidic platform for disparate density coculture reveals the dynamics of macrophage-mediated adipocyte clearance

**DOI:** 10.64898/2026.05.19.726422

**Authors:** Lim Yuan Bin, Kabigting Jessica Evangeline Tan, Cheam Mei Shan, Yusuke Toyama, Andrew W. Holle

**Affiliations:** Mechanobiology Institute, National University of Singapore, 117411, Singapore; Department of Biological Sciences, National University of Singapore, 117583, Singapore; Department of Biomedical Engineering, National University of Singapore, 117583, Singapore

## Abstract

Co-culturing cells with mismatched densities, where one cell type adheres to surfaces while the other floats, represents a fundamental challenge in cell biology. This is particularly evident in studying macrophage-adipocyte interactions, where macrophages must engage and clear lipid-rich apoptotic adipocytes, a process critical to understanding chronic inflammation in obesity and metabolic disease. The density disparity between macrophages, which sink and adhere to culture surfaces, and adipocytes, which float due to their lipid content, has prevented conventional co-culture approaches from achieving sustained cell-cell contact. To address this challenge, we developed a microfluidic system that confines adipocytes and lipid droplets in close proximity to macrophages. This platform features recessed micro-traps within the upper surface of a microfluidic chamber that trap buoyant objects while allowing media exchange and delivery of reagents for live-cell and immunofluorescence imaging. Time lapse imaging revealed that the dynamic process of macrophages-dead corpse interactions, showing that individual macrophages cannot engulf entire corpses but instead mechanically deform them. Furthermore, the platform successfully recapitulates the formation of Crown-Like Structures (CLS), clusters of macrophages surrounding dead adipocytes that are hallmarks of adipose tissue inflammation. Long-term culture revealed that CLS effectively clear lipids compared to partial macrophage engagement, providing mechanistic insights that were previously unattainable with standard histological approaches. Beyond the macrophage-lipid interaction, this platform has potential for studying interactions between adherent cells and buoyant targets, such as microplastics, opening new avenues for research where density mismatch poses a major barrier.

## Introduction

While adipose tissue has historically been viewed as an inert energy depot, it is now recognized as the body’s largest and most dynamic endocrine organ^1–4^, as it is essential for maintaining systemic lipid and glucose homeostasis^2,5,6^. While it is primarily comprised of adipocytes, it also contains a variety of stromal and immune cells^5,7,8^, most notably resident macrophages, which are critically important for clearing cellular debris and for their instructive role in orchestrating tissue development and remodelling^9–11^. Indeed, recent work has shown that specific macrophage subsets act as fundamental architects of adipose tissue, directing adipocyte stem cell fate and thereby controlling tissue plasticity during homeostasis^12^.

However, in conditions of metabolic stress, such as obesity and aging, these vital homeostatic and architectural functions are compromised^6,13,14^. High-lipid environments are known to directly alter the homeostatic phenotype of these resident macrophages, driving pathological outcomes^15^. Adipocytes become hypertrophic and die through inflammatory necrotic or pyroptotic pathways^16–18^. Instead of being neatly packaged for disposal, the cell ruptures, spilling a potent cocktail of damage-associated molecular patterns (DAMPs) and its massive lipid droplet into the extracellular space^19–21^. This event acts as a powerful “danger signal” that alters the local microenvironment. The dysregulation of such innate sensing pathways, where normally protective responses to the body’s own molecules become drivers of pathology, is a key feature of chronic inflammatory diseases^22^. If this lipid-rich debris is not cleared effectively, it triggers the chronic, low-grade inflammation that drives systemic metabolic dysfunction^19^.

While single macrophages are able to process most apoptotic cells on their own, they are incapable of engulfing a hypertrophic adipocyte or its massive lipid droplet^19,23^. When a macrophage encounters a target exceeding the critical size threshold for phagocytosis (∼25 µm in diameter), the clearance process necessitates a prolonged, cooperative effort from multiple cells^19^. This process results in the formation of “Crown-Like Structures” (CLS), which are assemblies of multiple macrophages physically surrounding a single dead adipocyte^16^. The abundance of CLS is strongly correlated with adiposity and insulin resistance, and for years they have been viewed as the epicentre of inflammation in obese adipose tissue^24,25^. However, little is known on their ability to impart force upon the apoptotic adipocytes that they collectively process, and the degree to which this mechanical aspect plays a role in clearance.

One of the major reasons that this knowledge gap persists is the inherent limitations of existing experimental platforms for studying adipocyte biology. *Ex vivo* approaches, such as tissue sectioning and explant imaging, provide important physiological context but are limited by their static nature or by poor imaging resolution due to light scattering^16,19^. Conventional *in vitro* models struggle with the fundamental density mismatch between adherent cells and buoyant lipids. While recent microfluidic approaches, such as hydrogel-embedded adipose-on-chip co-culture system^26^, provide valuable insights into tissue-level immune infiltration, their dense cellular packing and 3D matrices inherently limit the high-resolution optical tracking necessary to observe single cell-droplet mechanical interactions and leave lipids prone to fusion and lateral displacement, driven either by fluid shear during media exchange or by macrophage forces. Crucially, bulk co-culture platforms average cellular responses across heterogeneous populations, obscuring droplet-specific mechanical interactions and clearance kinetics.

Conversely, high-precision cell-pairing microfluidics enforce strict 1:1 confinement within narrow channels or microwells^27,28^. This structural bottleneck inhibits the free macrophage migration and collective engulfment required for Crown-like Structure (CLS) formation. Furthermore, mature adipocytes are orders of magnitude larger and more fragile than immune cells; single-trap architectures are difficult to optimise to simultaneously accommodate, flow, and stably isolate both cell types without inducing shear-induced rupture or losing buoyant targets to media exchange.

To address these technological gaps, we developed the Fluidic Adipocyte Trap for Tracking Immune Cell interactions (FATTIC). By pairing an open, height-optimized chamber with recessed ceiling micro-traps, this platform leverages buoyancy to spatially anchor individual targets while permitting free multi-cellular immune migration. Here, we detail the design, fabrication, and validation of this tool for studying complex macrophage interactions with large lipid droplets. We demonstrate that our device successfully recapitulates key *in vivo* phenomena, including the formation of CLS and the physical deformation of lipid droplets by macrophage-exerted forces, enabling the time-resolved quantification of clearance at a single-droplet level.

## Results and Discussion

### Stable, size-selective trapping of buoyant particles in a system that supports long-term cell culture

To address the challenge of studying interactions between adherent cells and buoyant targets, we designed and fabricated the Fluidic Adipocyte Trap for Tracking Immune Cell interactions (FATTIC) system. The device consists of a polydimethylsiloxane (PDMS) chip bonded to a glass coverslip, featuring a high-throughput array of recessed micro-traps (≥ 2,000 traps per chip) integrated into the ceiling of the microfluidic chamber (**Fig. 1A-C**). When lipid droplets are introduced into the chamber, they float up and are captured within the recessed traps, effectively immobilizing them against the ceiling (**Fig. 1B**). Light microscopy visualized the regularly spaced array of square traps (**Fig. 1C**), and confocal imaging of fluorescently labelled lipid droplets demonstrated successful capture within these structures (**Fig. 1D-E**).

**Figure 1:**
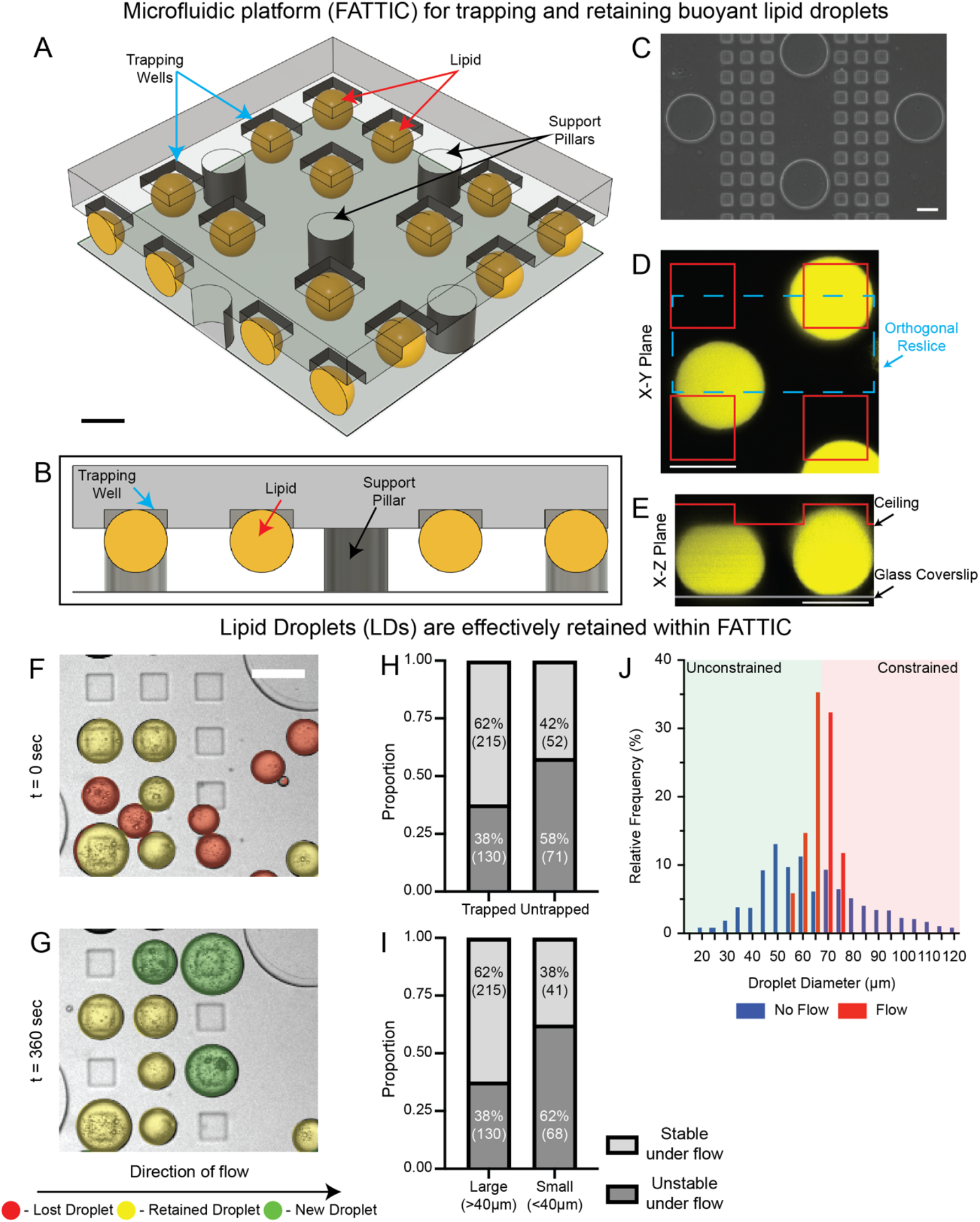
Design and validation of the microfluidic trap system for stable retention of buoyant particles. (A, B) Schematics illustrating device’s 3D architecture (A) and trapping principle (B), where buoyant objects are retained in recessed ceiling traps. Scale bar: 50 µm. (C) Light microscope image of the trap array. Scale bar: 100 µm. (D, E) Confocal fluorescence image (D) and corresponding cross-sectional view (E) showing a lipid droplet (yellow) secured within a trap (red outline), while an adjacent droplet remains un-trapped. Scale bar: 50 µm. (F, G) Brightfield images demonstrating droplet retention under fluid flow. Retained droplets are indicated with yellow circles, droplets that were washed away are indicated with red circles, and new droplets that entered the field of view are indicated with green circles. (H) Quantification of retention confirms that trap geometry, not just buoyancy, mediates stable retention, with a significantly higher percentage of trapped droplets remaining post-washout. (I) The system demonstrates inherent size-selectivity, as larger droplets (≥40 µm) are retained with significantly higher eWiciency under flow compared to smaller droplets. (J) Enrichment of lipid droplet size distribution through flow mediated washing. The initial heterogenous population in FATTIC (blue) is narrowed to a relevant range (red) by applying controlled flow, which washes away smaller droplets and enriches the trapped population, demonstrating the system’s inherent size-selective capability. Background shading represents the unconstrained droplets (green, droplets ≤65 µm, maintaining native spherical conformation) and the constrained droplets (red, droplets >65 µm, subjected to axial compression).

To validate the stability of this trapping mechanism against perturbations common in cell culture, such as media changes, we subjected the system to continuous fluid flow using a syringe pump (**Fig. S1A**). Brightfield imaging before and after the application of flow showed that droplets secured within the traps remained stably retained, while un-trapped droplets were washed away (**Fig. 1F-G**). Quantitative analysis confirmed that retention is mediated by the physical traps as a significantly higher percentage of trapped droplets remained post-washout compared to those in non-trap regions (**Fig. 1H**). We note that the observed retention rate (62% post-washout) reflects an engineering trade off which prioritises safe loading and high initial adipocyte viability over maximal trapping yield, as forcing fragile mature adipocytes into deeper traps in a narrower main chamber would induce shear rupture. The trap design also confers a size-selective capability to the system. By quantifying droplet retention under flow as a function of size, we observed that larger droplets (≥40 µm) were retained with significantly higher efficiency than smaller ones (**Fig. 1I**).

To demonstrate FATTIC’s capacity to reduce droplet heterogeneity and enrich the target population, we compared the initial loaded lipid droplets against the population retained following flow exposure (**Fig. 1K**). The pre-wash population exhibited a broad size distribution spanning 15 – 120 µm. Following flow induction, retained droplets shifted toward trap-compatible diameters, yielding a pronounced enrichment in the 55 – 75 µm window following image inspection. This size-selective retention arises from the trap geometry and controlled flow dynamics, which displace smaller droplets while stabilising larger targets. This retained droplet population aligns with the pathological observations that Crown-Like Structure formation predominantly occurs around hypertrophic adipocytes exceeding 50 µm^19^. Together, these results demonstrate that the system provides a robust and size-discriminating method for stably immobilizing buoyant particles, thereby establishing a controlled and reliable environment for subsequent cell-lipid interaction studies.

Crucially, the system is engineered to preserve the structural integrity of fragile adipocytes throughout the experiment. During initial cell loading, FATTIC’s shallow ceiling trap (15 µm) and deep microfluidic chamber (50 µm) prioritises cell viability by protecting large adipocytes from shear-rupture. For subsequent multi-day cell culture experiments, we developed an open-sided version of the chip that allows for passive nutrient and gas exchange (**Fig. S1B**). Using standard live/dead fluorescence assay, we confirmed that primary murine adipocytes remained viable within the device for 72 hours (**Fig. 2A,C**). Primary bone marrow-derived macrophages (BMDM) were also found to be viable in long-term culture (**Fig. 2B,D**), along with two standard cell lines (MDCK and MDA-MB-231) (**Fig. S2A**,**C**). The demonstrated long-term viability enables the observation of slow biological processes, such as CLS assembly and lipid digestion, that unfold over days.

**Figure 2:**
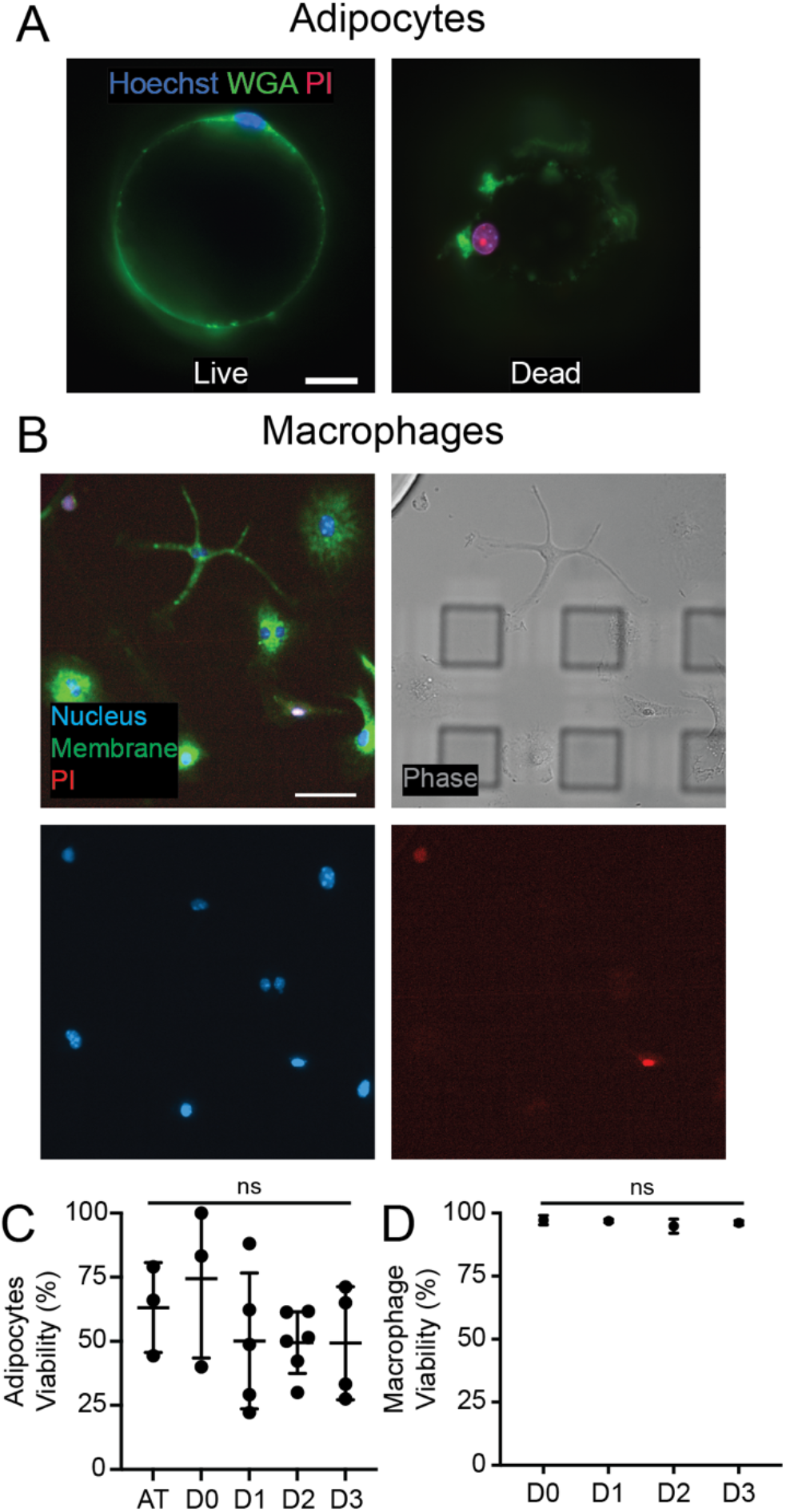
Biocompatibility and long-term cell viability. (A) Representative fluorescence images of live/dead staining in primary adipocytes. Live cells show an intact nucleus (Hoechst, blue) and plasma membrane (WGA, green), while dead cells lose membrane integrity and take up propidium iodide (PI, red). Scale bar: 10 µm. (B) Representative image of primary macrophages cultured in the system, stained as in (A). Scale bar: 50 µm. (C) Quantification of adipocyte viability, showing stable survival over 72 hours in the device. (D) Quantification of macrophage viability in monoculture over 72 hours. No significant loss of viability was observed, demonstrating the system’s biocompatibility for long-term macrophage culture.

### Dynamic observations of direct macrophage-lipid interactions

In obesity, hypertrophic adipocytes frequently undergo pyroptosis or necrosis, leading to plasma membrane rupture^16^. Consequently, the target encountered by macrophages *in vivo* is often a cell-free lipid remnant. Yet, the specific biophysical mechanisms by which macrophages engage with and clear this exposed lipid without the scaffold of an intact membrane remain poorly understood. To address this, we utilised FATTIC to model this terminal phase of clearance, exploring the direct interactions between adherent BMDMs and buoyant, adipocyte-derived lipid droplets.

We first verified that FATTIC supports the extended co-culture necessary to observe these prolonged interactions. Both macrophages and other adherent cell lines (MDCK and MDA-MB-231) were found to remain viable for over 72 hours in the presence of lipid droplets (**Fig. 3A-B, Fig. S2B**,**D**). A critical design feature of this co-culture model is that buoyant lipid droplets are stably anchored in recessed ceiling traps while the chamber floor remains laterally unconstrained which permits unrestricted macrophage migration and collective assembly around the droplets.

**Figure 3:**
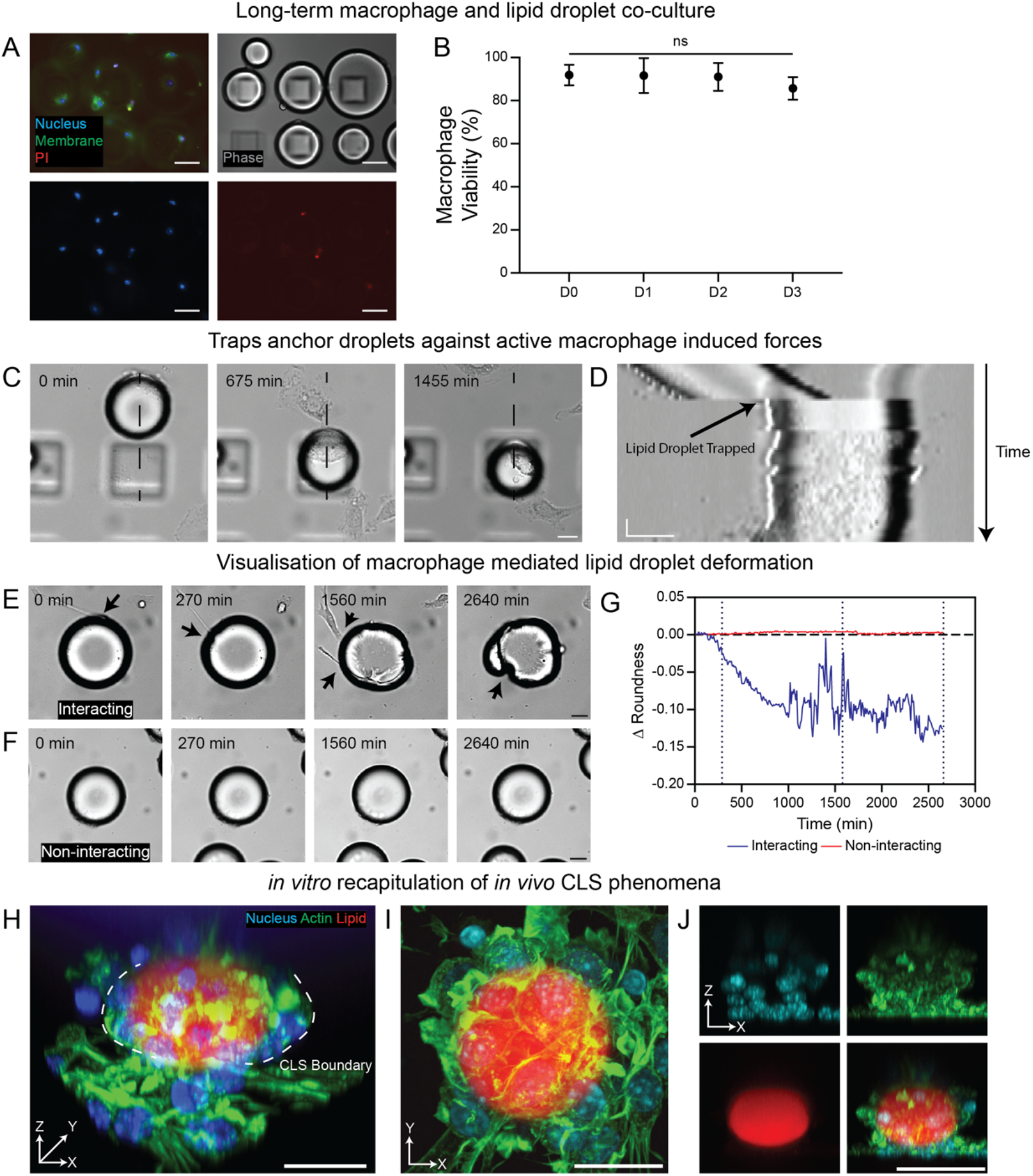
Visualization of key macrophage-lipid droplet interactions, including physical deformation and crown-like structure (CLS) formation. (A) Representative fluorescence image of primary macrophages co-cultured with murine-derived lipid droplets. Cells were stained with Hoechst (Blue), WGA (Green), and PI (Red). Scale bar: 50 µm. (B) Macrophage viability remained high over 72 hours in co-culture, confirming the system’s biocompatibility. (C, D) Validation of mechanical stability against cellular forces. Representative time-lapse brightfield images (C) showing a lipid droplet being actively pushed by macrophages but successfully entering and retaining within the trap. Corresponding kymograph (D) illustrates the stable long-term immobilization of the droplet after capture, despite continuous macrophage contact. Scale bar: 20 µm (Horizontal), 120 min (Vertical). (E) Time-lapse imaging captures the physical deformation of a lipid droplet by encroaching macrophages (arrows). Scale bar: 20 µm. (F) As a negative control, time-lapse imaging of a lipid droplet in the absence of macrophages shows no significant change in shape over the same period. Scale bar: 20 µm. (G) This visual observation is confirmed by roundness analysis, which quantifies significant shape fluctuations only in lipid droplets that are in direct contact with macrophages, as compared to lipid droplets alone. For temporal reference, vertical dotted lines on the graph correspond to the timepoints shown in (E, F) (0, 290, 1580, and 2660 min), confirming active mechanical engagement by macrophages. (H - J) A representative CLS formed by macrophages surrounding a lipid droplet. Cells were stained with Hoechst (Blue), Actin (Green), and lipid droplets were stained with LipidTox (Red). 3D reconstruction (H), Maximum intensity projection (MIP) (I) and orthogonal (XZ) (J) views demonstrate the 3D envelopment of the droplet by the cells. Scale bar: 50 µm.

Indeed, we observed that actively migrating macrophages were physically displacing untrapped lipid droplets (**Fig. S3**). However, trapped droplets were able to successfully resist these macrophage-induced movements (**Fig. 3C-D**), allowing for long-term tracking of macrophage-lipid interactions in the same field of view.

Continuous imaging revealed that these interactions included dramatic deformations of lipid droplets in contact with macrophages (**Fig. 3E**), while droplets not in contact with macrophages remained spherical (**Fig. 3G**). In addition to single macrophage interactions, the formation of CLS was observed, in which multiple macrophages migrated towards the droplet and fully enveloped it, similar to what is observed in CLS *in vivo* (**Fig. 3H-J**). Capturing this dynamic multicellular coordination is a critical capability of the FATTIC system enabled by the free movement of macrophages in the microfluidic chamber. Having successfully captured CLS formation, we next sought to determine whether CLS formation is necessary for lipid clearance.

### Multicellular macrophage organisation is necessary for lipid clearance

The degree to which macrophages are capable of processing lipid droplets is difficult to quantify in *in vivo* or *ex vivo* systems. To measure this in our *in vitro* system, we co-cultured RAW 264.7 macrophages with fluorescently-labelled lipid droplets and monitored their interactions over 48 hours (**Fig. 4A**). We categorized these interactions based on the degree of macrophage envelopment, non-CLS engagement (< 75% target coverage) and CLS formation (≥ 75% target coverage).

**Figure 4:**
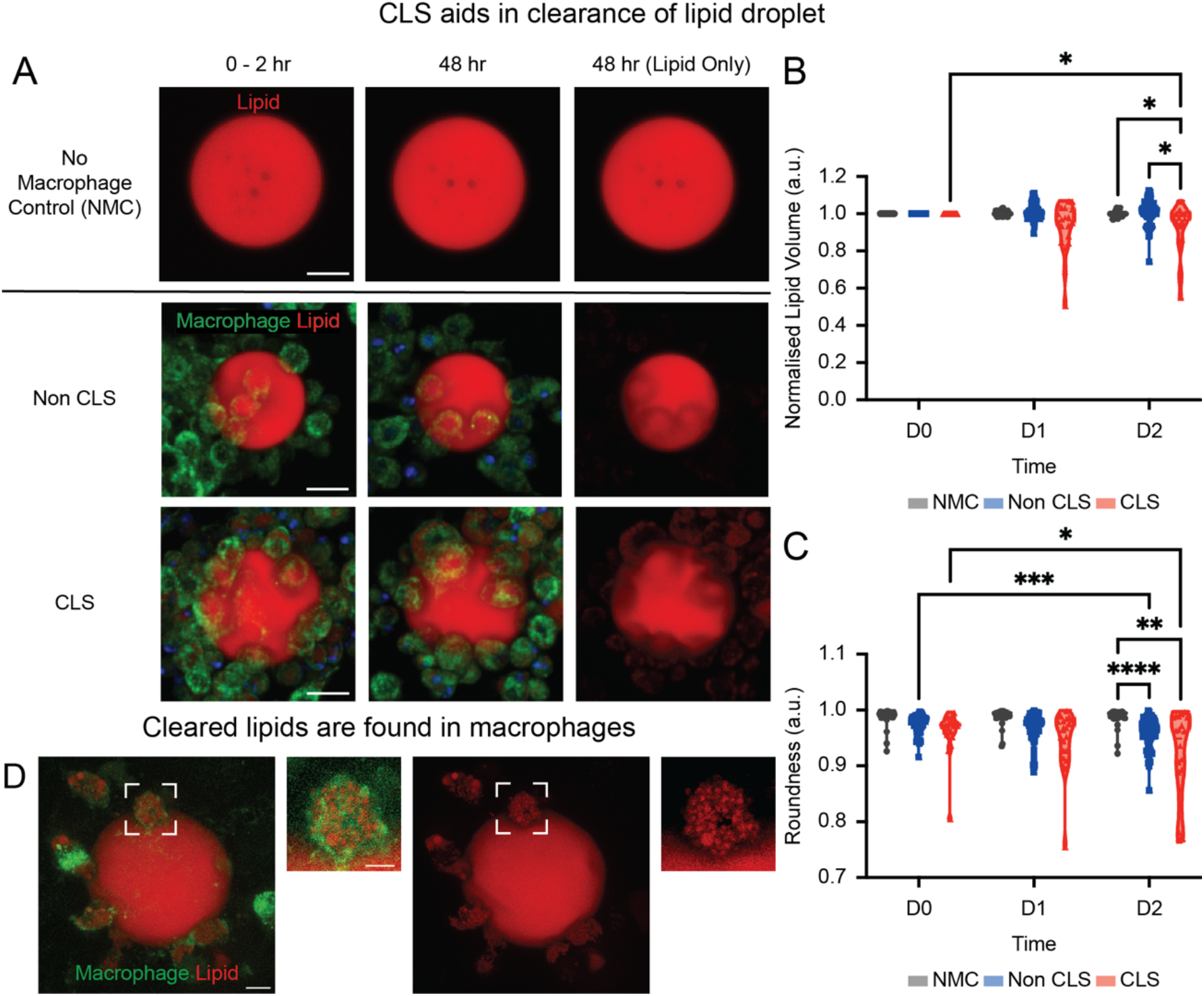
Macrophage-mediated lipid clearance is enhanced by Crown-like Structure (CLS) formation. (A) Representative images of lipid droplets (LipidTox, red) co-cultured with RAW 264.7 macrophages (WGA membrane stain, green; Lipid: LipidTox, Red) or no-macrophage control at 0-2 hours and 48 hours. Scale Bar: 20 µm. (B) Quantification of normalised extracellular lipid droplet volume over time. Individual droplet measurements are aggregated by time to show distribution and trend. ANOVA reveals a significant reduction in lipid volume for the CLS condition compared to Non-CLS and NMC controls at Day 2. (C) Quantification of lipid droplet roundness over time. ANOVA demonstrates a significant decrease in roundness for macrophage-interacting droplets compared to sable controls at Day 2. (D) Fluorescence images showing intracellular lipid droplets (BODIPY, red) within primary macrophages (WGA, green) after 7 days of co-culture, confirming that extracellular lipid reduction results from cellular uptake. Scale bars: 10 µm (main), 5 µm (inset). *p < 0.05, **p < 0.01, ***p < 0.001, ****p < 0.0001

Lipid droplets in cell-free control (NMC) samples did not show any significant loss of volume of 48 hours, showing that any lipid loss must be a consequence of macrophage interactions (**Fig. 4B**). Crucially, Non-CLS droplets also demonstrated no significant volume loss, indicating that simple physical contact is insufficient for lipid clearance. In contrast, CLS droplets exhibited a significant, time-dependent reduction in volume, indicating that CLS formation drives effective lipid clearance compared to limited macrophage engagement. Concurrently, CLS droplets displayed a marked decrease in roundness, reflecting progressive mechanical deformation (**Fig. 4C**). Statistical analysis using a two-way ANOVA with Tukey’s post-hoc test confirmed that clearance kinetics and morphological changes were significantly accelerated only in the CLS condition.

Finally, to identify the destination of processed lipid, we performed a separate long-term coculture using primary macrophages. After seven days, high resolution microscopy showed the emergence and massive accumulation of intracellular lipid droplets within the BMDMs, revealing that the surrounding cells were internalizing the lipids (**Fig. 4D**).

Collectively, these findings confirm that the extracellular volume loss observed during CLS envelopment is driven by macrophage lipid internalisation. By successfully capturing CLS progression from initial multi-cellular recruitment to physical deformation to foam cell formation, we demonstrate that FATTIC bridges a critical gap in modelling adipose tissue homeostasis.

## Conclusion

A wide variety of interactions occur *in vivo* between objects with disparate densities, including immune cells and microplastics, gas exchange at the air-liquid interface, and macrophage scavenging of apoptotic adipocytes. However, due to the open nature of the majority of cell culture tools, controlled analysis of objects with disparate densities is challenging *in vitro*. Here, we have established a platform capable of long-term high-resolution tracking of mechanical deformation in density-mismatched cells and lipid droplets. This system revealed for the first time that individual macrophages are capable of dynamically deforming bare lipid droplets, and that lipid processing is a collective process that requires complete envelopment of the lipid droplet by groups of macrophages, reminiscent of the Crown-Like Structures (CLS) observed *in vivo*. Unlike conventional co-culture or cell-pairing platforms, FATTIC uniquely couples stable single-droplet immobilisation with physiologically relevant multi-cellular dynamics. The establishment of this system has the potential to play a major role in advancing our understanding of cell-lipid interactions, both by allowing for a more complete characterization of the specific identities of lipids that macrophages are most efficient at processing and by providing a platform for high-throughput investigations of biochemical mediators of macrophage-lipid interactions.

## Author contributions

Y.T. and A.W.H. conceived the study and designed research. L.Y.B, Y.T. and A.W.H. designed the experiments and methods for data analysis. L.Y.B performed the microfluidic and biological experiments with assistance from C.M.S. and K.J.E.T., who assisted with murine tissue harvesting and primary cell isolation. L.Y.B, Y.T. and A.W.H. analysed the data. L.Y.B, Y.T. and A.W.H. wrote the manuscript. Y.T. and A.W.H. supervised the project. All authors read and commented on the manuscript.

## Data Availability

The data supporting this article, including the microfluidic device design files, the source data for all quantitative figures, and the Python scripts used for Linear Mixed Effects Model analysis are available in the Zenodo repository at h-ps://doi.org/10.5281/zenodo.20151348.

## Acknowledgements

The authors acknowledge the support of the National Research Foundation of Singapore under an NRF Fellowship to AWH (No. NRFF13-2021-0114). This work is also supported by a Singapore Ministry of Education Tier 2 grant (T2EP30220-0033 to Y.T.) and by the National Research Foundation of Singapore under its mid-sized grant (NRF-MSG-2023-0001 to Y.T. and AWH). We thank Gianluca Grenci, Mona Suryana, and Sebastian Foo Junwei at the Nano and Micro Fabrication Core Facility of the Mechanobiology Institute (MBI) for their expert assistance with device fabrication. We also thank Vinod S/O Prabhakaran, Nicole Lee Jia Wen, and Li Yixuan from MBI for providing the MDCK and MDA-MB-231 cell lines used in viability assays. We are grateful to Rong Li and their lab members Lan Xi and Shen Xingyu, as well as Chii Jou Chan and their lab member Apoorva Shivankar for facilitating access to murine tissues through the MBI tissue-sharing program. We also thank Gao Xu for technical training and advice regarding PDMS device bonding. We also thank the staff at the MBI Wet Lab Core for infrastructural support, and Hui Ting Ong and Fei Li Jasmine Chin at the MBI Microscopy Core for their support with microscopy.

## Experimental

### Animals

Mice were obtained through an institutional tissue-sharing program, where tissues were collected post-experiment from unrelated studies conducted by other researchers within the institute. This approach minimized animal use and promoted resource sharing. The animals were maintained under a 12-hour light/dark cycle with ad libitum access to food and water. They were fed a standard chow diet consisting of 7% simple sugars, 3% fat, 50% polysaccharides, and 15% protein (w/w). Euthanasia was performed using CO_2_ inhalation by personnel certified and approved under the Institutional Animal Care and Use Committee (IACUC). All animal procedures complied strictly with IACUC-approved protocols.

### Cell Culture and Primary Cell Isolation

RAW 264.7 macrophages (TIB-71, ATCC) were cultured in RPMI 1640 medium supplemented with 10% fetal bovine serum (FBS; 10500-064, Thermo Fisher Scientific), 1% penicillin-streptomycin (P/S; 15140122, Thermo Fisher Scientific). MDA-MB-231 and MDCK were cultured in Dulbecco’s Modified Eagle’s Medium (DMEM; 11995073, Thermo Fisher Scientific) supplemented with 10% fetal bovine serum (FBS; 10500-064, Thermo Fisher Scientific), 1% penicillin-streptomycin (P/S; 15140122, Thermo Fisher Scientific). Cells were maintained at 37°C in a humidified incubator with 5% CO_2_.

The isolation of bone marrow-derived macrophages (BMDMs) was performed following established protocol^29^. Briefly, femurs and tibias were dissected from mice using forceps and scissors, ensuring the removal of all muscle and fibrous tissue. If necessary, the bones were wiped with a Kimwipe moistened with 70% ethanol to eliminate residual fibres. For bone marrow extraction, two Eppendorf tubes were prepared: a 1.5 mL tube (outer tube) and a 0.5 mL tube (inner tube) with a pre-pierced hole at the bottom. Both ends of the femurs and tibias were cut, and the bones were placed into the inner tube. Centrifugation at 12,000 x g for 30 seconds flushed the bone marrow into the larger tube while retaining the bones in the smaller tube. Erythrocytes were removed by resuspending the bone marrow in 1X RBC lysis buffer, incubating for 5 minutes, and centrifuging at 300 x g for 5 minutes. This step was repeated as needed to ensure thorough removal of erythrocytes. The resulting bone marrow was resuspended in complete RPMI 1640 medium supplemented with 10% fetal bovine serum (FBS; 10500-064, Thermo Fisher Scientific), 1% penicillin-streptomycin (P/S; 15140122, Thermo Fisher Scientific), and 2 ng/mL macrophage colony-stimulating factor (M-CSF; M6518, Sigma-Aldrich). The bone marrow suspension was cultured overnight in T25 tissue culture flasks. The following day, 500 µL of media containing floating myeloid progenitor cells was collected and seeded onto non-tissue culture-treated petri dishes containing bone marrow differentiation medium. Cultures were maintained for 7 days, with the medium replenished every 3–4 days. By the end of the 7-day culture period, differentiated macrophages were ready for downstream applications.

Adipose tissue was obtained and processed according to the protocol outlined previously^30^. Adipose tissue was harvested from mice following euthanasia and processed for the isolation of mature adipocytes. The abdominal region was sterilized with 70% ethanol, and an incision was made to expose the perigonadal white adipose tissue (pWAT). The desired adipose depots were carefully excised and transferred to petri dishes containing phosphate-buffered saline (PBS) supplemented with 1% penicillin/streptomycin (P/S). To preserve tissue viability, processing occurred within one hour of collection. The adipose tissue was weighed and minced into small fragments (∼1–2 mm^3^) using sterile scissors in PBS + P/S. The minced tissue was washed three times with fresh PBS + P/S, transferring the slurry to a new sterile petri dish after each wash to remove impurities. The tissue was then transferred into a 50 mL Falcon tube containing a collagenase solution (PBS with 1% bovine serum albumin [BSA] and 0.2% collagenase, prepared at 10 mL per 2 g of tissue). The tube was incubated at 37°C on an orbital shaker set to 200 rpm for 30 minutes. After enzymatic digestion, the tissue suspension was centrifuged at 300 × g for 5 minutes at 22°C. The tube was inverted to mix the contents and centrifuged again under the same conditions. Mature adipocytes, which floated to the top of the solution, were carefully aspirated using a Pasteur pipette and transferred to a new 15 mL Falcon tube. The adipocytes were washed with 8 mL of PBS + P/S to remove residual impurities and centrifuged again at 300 × g for 5 minutes at 22°C. The purified adipocytes were gently collected using a pipette tip and transferred to sterile Eppendorf tubes. For downstream applications, adipocytes were resuspended in DMEM/F12 medium supplemented with 10% fetal bovine serum (FBS) and 1% P/S.

### Isolation and generation of lipid droplets from murine adipocytes

Adipocytes from the previously isolated layer were transferred into a new tube and exposed to ultraviolet (UV) light for one hour to induce cell death. Following this treatment, the suspension was centrifuged again at 5,000 rpm for 5 minutes at room temperature, resulting in two distinct layers: the adipocyte and lipid layer. Lipids from 100 µL of the irradiated adipocyte layer were resuspended in 900 µL of media and vortexed at maximum speed for 30 seconds to generate a lipid droplet emulsion. These lipid droplets were then used for downstream applications.

### Fabrication of FATTIC chips

Silicon master moulds were fabricated using a two-step photolithography process. Polydimethylsiloxane (PDMS) was prepared at a 10:1 base-to-crosslinker ratio (SYLGARD Silicone Elastomer), poured onto the silicon mould, degassed for 5 min, and cured at 80°C for two hours. After curing, the PDMS chips were carefully cut from the wafer. Two distinct chip designs were fabricated to suit different experimental needs. For lipid washout experiments, a “closed” design was created using rectangular chips with two 0.5 mm biopsy punches for an inlet and outlet (**Fig. S1A**). For cell culture and co-culture experiments, an “open” design was used, featuring openings on all four sides and a central hole for cell seeding (**Fig. S1B**). Both PDMS chips and glass coverslips were treated with oxygen plasma and irreversibly attached to form microfluidic devices. The internal surfaces of the bonded chips were then coated with rat tail collagen I dissolved in 0.1% acetic acid solution at a concentration of 150 µg/mL (A1048301, Gibco). After incubation at 37°C overnight, the chip was rinsed 3 times in PBS and UV disinfected for 30 min.

### Cell seeding in FATTIC chips

The following procedure was used for all cell types (BMDMs, RAW 264.7, MDA-MB-231, and MDCK). Cells were trypsinized, and a 20 µL suspension containing approximately 20,000 cells was carefully added to the central hole of the open chip design. The chip’s design allowed the centre and four sides to remain exposed to the atmosphere, while the sub-chip space, approximately 65 µm in height, does not restrict cell movement. Within 30 minutes, cells adhere to the collagen-coated coverslip, after which the well was filled with media and incubated overnight. For adipocyte seeding, live adipocytes were prepared by resuspending 100 µL of isolated adipocytes in 900 µL of media. 20 µL of the suspension was pipetted and slowly injected into the FATTIC chip. Excess adipocytes outside the chip were removed by washing the system three times with 1X PBS. To seed lipid droplets, 20 µL of the prepared murine lipid droplet suspension was pipetted and slowly injected into the FATTIC chip. The chip’s height of approximately 65 µm allowed for the selective trapping of lipid droplets 65 µm or larger in diameter, while smaller droplets were flushed out. Excess lipid droplets outside the chip were removed by washing the system three times with 1X PBS. Following overnight incubation, the media was aspirated and lipid droplets were added to the chip as described above. After PBS washes via gravitational flow, complete media was added to submerge the chip.

### Lipid droplet washout

The FATTIC chip containing lipid droplets was connected to a syringe pump via tubing attached to the entry port and linked to a waste container at the exit port. Sequential flows of 25 µL, 50 µL, 100 µL, 200 µL, and 400 µL per min were introduced into each chip. Before imaging, lipid droplets were stained with BODIPY™ 493/503 (D3922, Invitrogen) for 30 minutes. Timelapse imaging was performed using a Nikon A1R MP laser scanning confocal microscope equipped with a Nikon 10x objective lens and controlled via NIS-Elements software. Imaging occurred at 1-second intervals, with flow introduced 30 seconds after imaging began, maintained for 3 minutes, and ceased 30 seconds before the end of recording. Static images were also captured before and after flow to evaluate lipid trapping efficiency within the chip. For quantification, droplets were classified as either ‘Trapped’ or ‘Untrapped’ based on their location relative to the ceiling traps. Retention was defined by whether a droplet remained in its original Region of Interest (ROI) post-washout; displacement from the ROI was classified as “unstable under flow”. The initial size distribution of the murine lipid droplets in FATTIC was measured via brightfield imaging (droplets ≥120 µm were excluded). Following flow induction, retained droplets were imaged and measured to determine the post-washout size distribution, with final droplet inclusion confirmed with image inspection to ensure consistent morphology and reliable tracking.

### Tracking Cell Survival in FATTIC

At 2, 24, 48, and 72-hour timepoints, chips were removed from the incubator, and Hoechst (1:1000 dilution factor [DF]), WGA Oregon Green (1:100 DF), and propidium iodide (PI, 1:1000 DF) were added to the chips. After a 30-minute incubation, the dyes were aspirated, and the chips were washed three times with 1X PBS. Imaging was performed on a CD7 epifluorescence microscope. Image analysis was conducted using Fiji (ImageJ) across multiple randomly selected fields of view. The total number of cells was determined by counting Hoechst-positive nuclei, and dead cells were identified as PI-positive nuclei. Viability was calculated as the percentage of PI-negative cells relative to the total number of cells.

### Crown-like structure (CLS) Formation and Visualisation

To induce the formation of robust CLS suitable for high-resolution imaging, an increased RAW 264.7 macrophage seeding density was utilized. A 20 µL suspension containing approximately 100,000 cells was carefully added to the central port of the FATTIC chip and co-cultured with lipid droplets. Following a co-culture period of four days, samples were fixed for 15 minutes at room temperature using 4% paraformaldehyde (PFA, Cat# 50-980-487, Fisher Scientific). Subsequently, the chips were incubated with a staining cocktail containing Hoechst 33342 (1:1000 dilution) to label nuclei, Phalloidin AF488 (1:500 DF, A12379, ThermoFisher Scientific) to visualize the F-actin cytoskeleton, and LipidTox Deep Red (1:1000 DF, H34477, ThermoFisher Scientific) to stain the lipid droplets. High-resolution, three-dimensional images of the resulting CLS were then acquired via BC43 confocal microscopy.

### Assessment of Trap Mechanical Stability

To validate the ability of the FATTIC traps to retain lipid droplets against macrophage forces, time-lapse imaging was performed. Co-cultures were imaged using brightfield microscopy at 10 to 15 minute intervals. Kymographs were generated in Fiji (ImageJ) by defining a linear region of interest (ROI) across the droplet and/or trap axis to visualise and verify droplet immobilisation over time.

### Quantification of lipid droplet clearance

To quantitatively assess the rate of macrophage-mediated lipid clearance, the shape and size of individual lipid droplets were tracked over time in both co-culture and no-macrophage control (NMC) conditions. For imaging, co-cultures were stained for 1 hour with Hoechst 33342 (1:1000), WGA Oregon Green (1:100), and LipidTox Deep Red (1:500). Following incubation, the staining solutions were aspirated and the chips were washed three times with 1X PBS. Live-cell imaging was performed at 2, 24, and 48 hour timepoints using a BC43 spinning disk confocal microscope, with the LipidTox channel providing high contrast for automated analysis. These images were analyzed in Fiji (ImageJ) by applying an intensity threshold to generate a segmentation mask of the droplet’s boundary (**Fig. S4**). To assess clearance, a perfect circle was algorithmically fitted to the mask to calculate the 2D radius, from which lipid droplet volume was estimated assuming a spherical geometry. Because this 2D-to-3D extrapolation introduces variances during non-spherical, macrophage-induced deformation, transient volume fluctuations were anticipated. Therefore, to ensure robust quantification, volume measurements were normalised to the mean volume of the No Macrophage Control (NMC) group at the initial time point. To assess deformation, droplet roundness was measured using Fiji’s built in shape descriptors. For analysis, droplets were classified into three conditions: No Macrophage Control (NMC), Non-CLS (<75% macrophage coverage), and Crown-like Structures (CLS, ≥75% macrophage coverage).

### Statistical Analysis

Statistical analyses were performed using GraphPad Prism (version 11.0). Data were assessed for normal distribution and homogeneity of variance prior to testing. For experiments evaluating two independent variables, such as the lipid clearance and deformation kinetics of stationary, trapped droplets (Fig. 4B, C), a two-way analysis of variance (ANOVA) was performed, followed by Tukey’s post-hoc test for multiple comparisons. Non-parametric comparisons between two groups (Supplementary Fig. 2C, D) were conducted using the Mann-Whitney *U* test. For non-parametric analyses involving three or more groups, including cell viability assessments (Fig. 2C, 2D, 3B), the Kruskal-Wallis test was applied, followed by Dunn’s multiple comparisons test. Statistical significance was defined as p < 0.05.

### Visualization of Intracellular Lipid Droplets

To confirm that the clearance of extracellular lipid droplets was due to cellular uptake, the accumulation of intracellular lipids within macrophages was visualized after long-term co-culture. Primary BMDMs were co-cultured with lipid droplets in the FATTIC system for seven days with daily media replenishment. Following this co-culture period, the cells were stained for 1 hour with BODIPY 493/503 (1:1000) to label lipids and WGA Texas Red (1:100) to delineate the macrophage plasma membrane. After incubation, the dyes were aspirated, and the chips were washed three times with 1X PBS. High-resolution confocal Z-stacks were then acquired using a Nikon A1R confocal microscope with a 100x objective to visualize the accumulation of distinct, intracellular lipid puncta within the macrophage cytoplasm, confirming cellular uptake and foam cell formation.

