## Supplementary Figures 1-4 for "A confining microfluidic platform for disparate density coculture reveals the dynamics of macrophage-mediated adipocyte clearance"

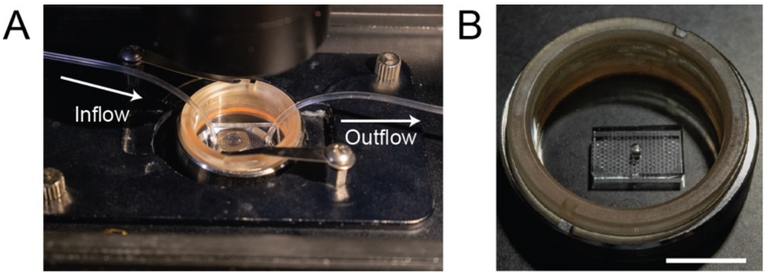


***Supplementary Figure 1*: *Photograph of experimental setup.*** *(A) Assembled "closed" FATTIC chip for washout experiments featuring inflow and outflow ports. (B) The "open" chip designed for long-term cell culture. Scale Bar: 1 cm.*

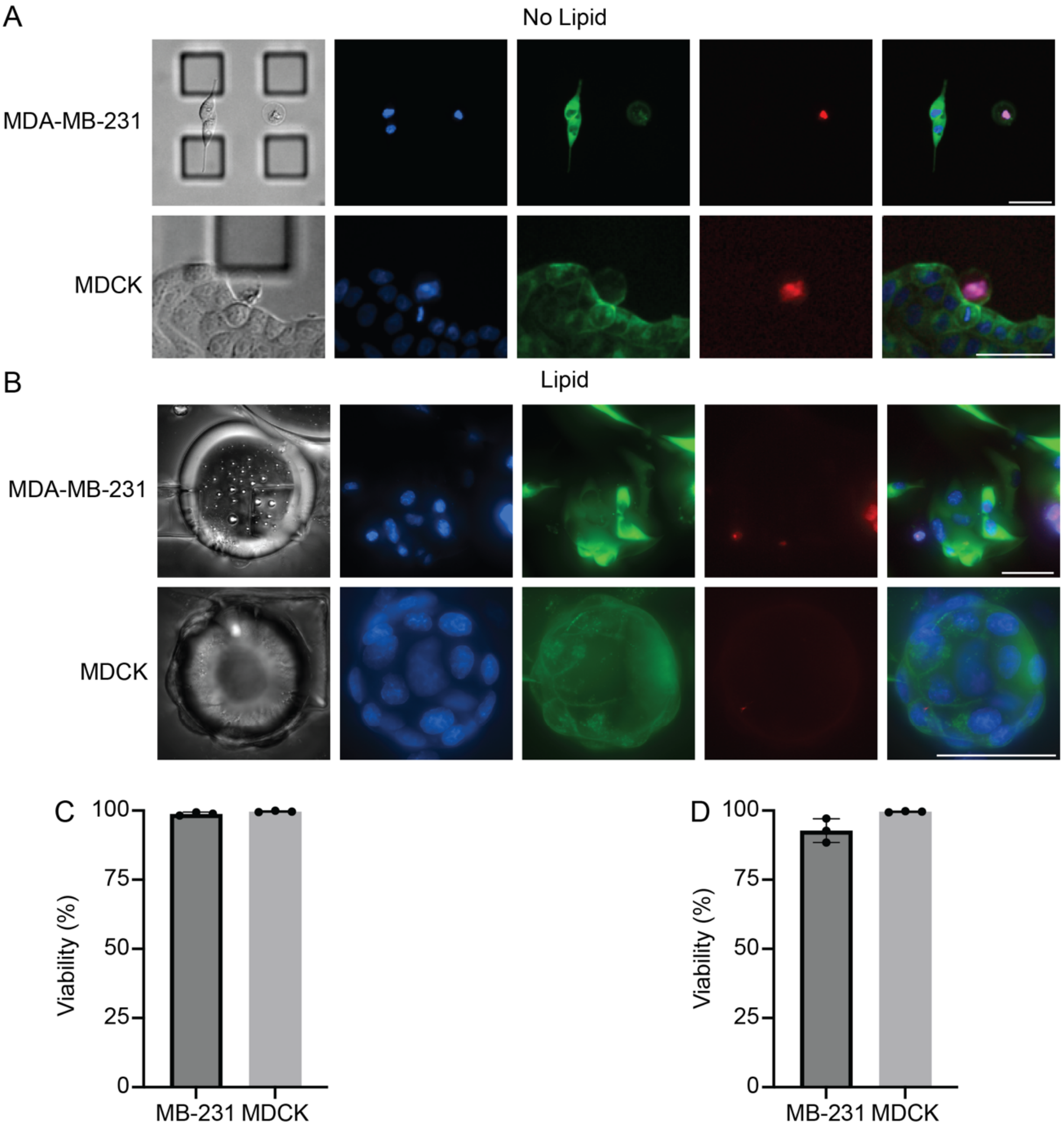


***Supplementary Figure 2**: **Long-term survival of different adherent cell lines** (A, B): Representative fluorescence images of MDA-MB-231 and MDCK cells cultured for 72 hours in the FATTIC system. Cells were cultured in standard medium (A) or in co-culture with a high concentration of murine-derived lipid droplets (B). Images show high cell viability in both conditions, with live cells indicated by intact membranes (WGA, green) and propidium iodide (PI)-negative nuclei, and rare dead cells marked by PI-positive nuclei (red). All nuclei are stained with Hoechst (blue). Scale bar = 50 µm. (C, D): Quantification of cell viability over 72 hours for MDA-MB-231 and MDCK cells cultured without lipids (C) and with lipids (D). Data are presented as mean ± SD from n=3 independent experiments.*

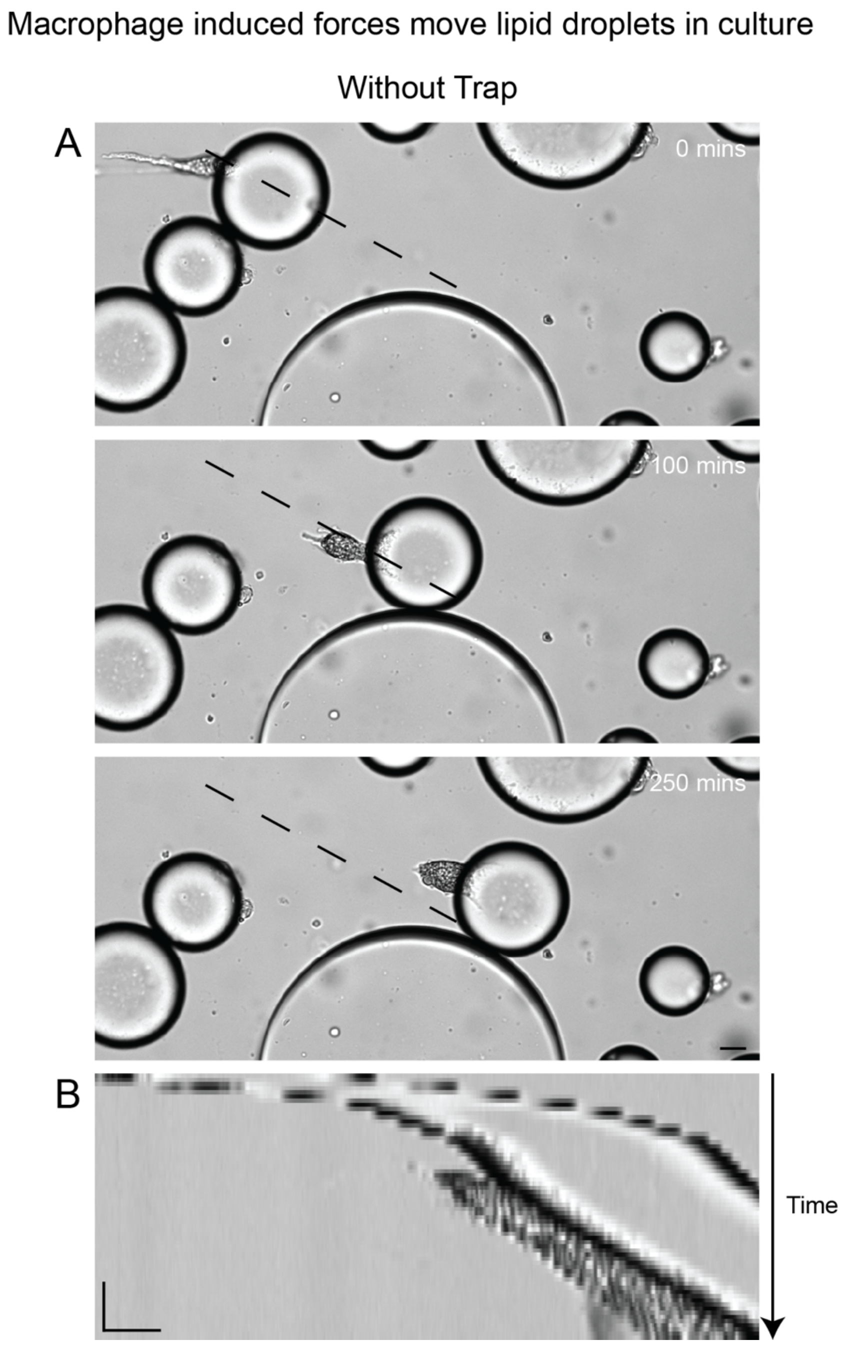


***Supplementary Figure 3*: *Macrophages actively displace lipid droplets in the absence of microfluidic traps.*** *(A) Representative time-lapse brightfield images showing a lipid droplet actively pushed and displaced across the field of view by the forces of interacting macrophages. (B) Corresponding kymograph of the droplet in (A), illustrating significant spatial drift over time. Scale Bar: 20 µm (A), 20 µm (horizontal, 120 min (vertical) (B).*

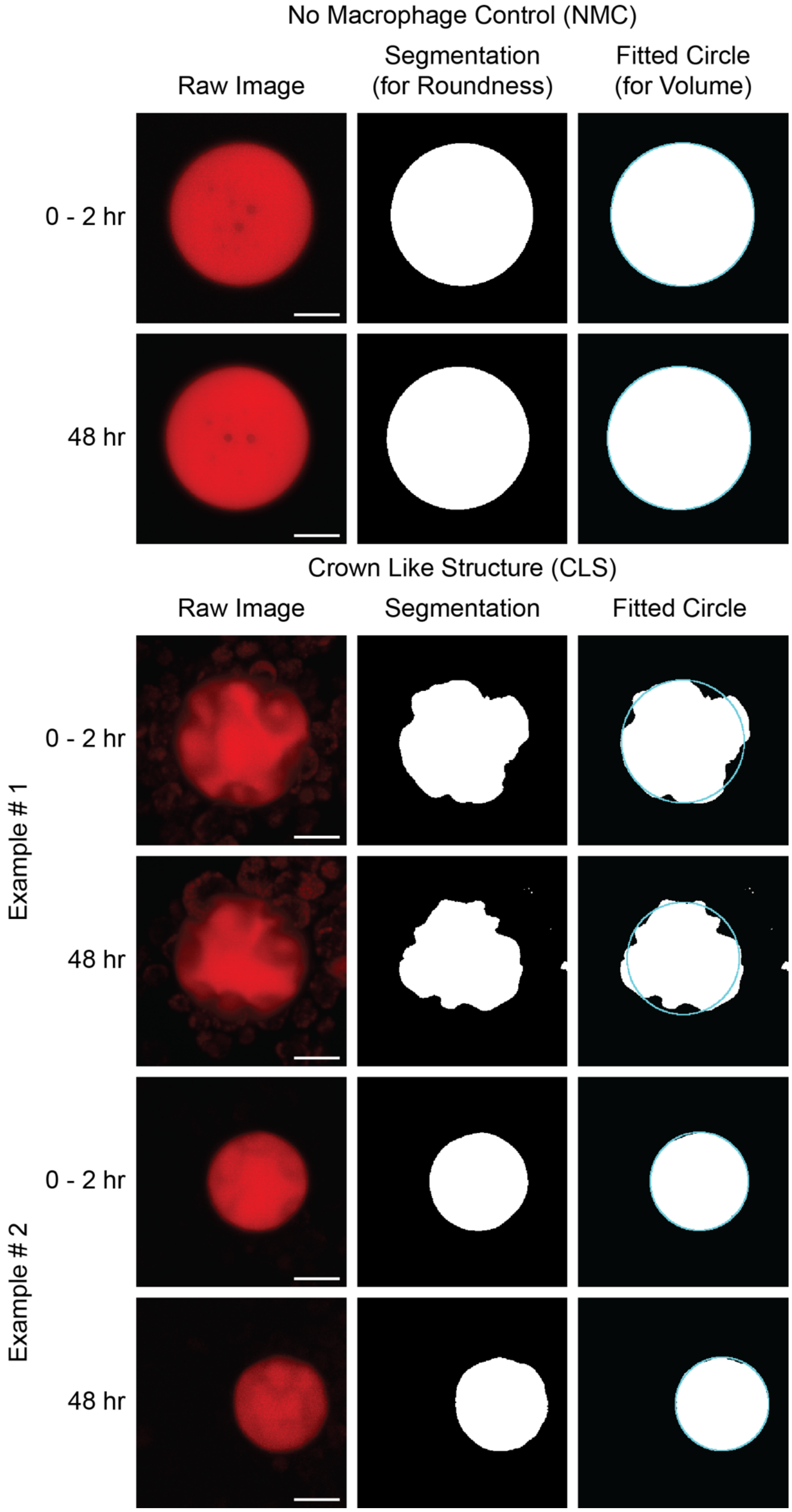


***Supplementary Figure 4: Demonstration of segmentation behaviour on lipid droplets over a 48-hour time course.*** *The fitted circle (light blue) highlights deviations between the 2D segmentation mask and the true droplet*

*boundary, particularly under active macrophage-induced deformation (CLS). This illustrates the inherent challenge of estimating volume from 2D projections of 3D deforming droplets, rationalizing the observed measurement noise in the main text.*
